# Image-based quantification of histological features as a function of spatial location using the Tissue Positioning System

**DOI:** 10.1101/2022.10.12.511979

**Authors:** Yunguan Wang, Ruichen Rong, Yonglong Wei, Tao Wang, Guanghua Xiao, Hao Zhu

## Abstract

Tissues such as the liver lobule, kidney nephron, and intestinal gland exhibit intricate patterns of zonated gene expression corresponding to distinct cell types and functions. To quantitatively understand zonation, it would be important to measure cellular or genetic features as a function of position along a zonal axis. While it is possible to manually count, characterize, and locate features in relation to the zonal axis, it is very difficult to do this for more than a few hundred instances. We addressed this challenge by developing a deep-learning-based quantification method called the “Tissue Positioning System” (TPS), which can automatically analyze zonation in the liver lobule as a model system. By using algorithms that identified vessels, classified vessels, and segmented zones based on the relative position along the portal vein to central vein axis, TPS was able to spatially quantify gene expression in mice with zone specific reporters. TPS could discern expression differences between zonal reporter strains, ages, and disease states. TPS could also reveal the zonal distribution of cells previously thought to be randomly distributed. The design principles of TPS could be generalized to other tissues to explore the biology of zonation.

The software is available at https://github.com/yunguan-wang/Tissue_positioning_system.

## Introduction

Tissues such as the liver, kidney, and intestine carry out diverse biological functions. An elegant division of labor among cells within these tissues is spatially organized into “zones” that express specialized genetic programs ^1^. For example, different zones in the liver carry out distinct metabolic functions. Liver zonation is dynamic and can change during development, aging, and disease, making it important to measure changes in zonal boundaries and gene expression patterns under different biological conditions. Necessary for this kind of analysis is the quantification of important features within a tissue and understanding how they are spatially organized. For any given coordinate on an image, one would like to know the position with respect to anatomical landmarks that define the zonal axes. Once this “location” is obtained, phenotypes and gene expression at this position can be assessed as a function of anatomical coordinates, or vice-versa. This kind of analysis is ordinarily performed in a qualitative fashion by visual inspection of histology images. However, it would be preferable to obtain these data using the systematic and quantitative evaluation of images. Dedicated computational tools to parse zonation in an automated fashion have not been developed. The liver is the ideal setting to establish an algorithm to measure tissue features as a function of spatial position.

The lobule is the basic functional unit of liver tissue that is shaped like a submarine in three dimensions, hexagonal when cross-sectioned, and repeated throughout the liver^1,2^. Lobules are outlined by portal veins (PVs) at the vertices and one central vein (CV) in the center. Portal venous blood from the intestine intermixes with oxygenated arterial blood in the portal region, then flows through the sinusoids and collects in the CVs before returning to the heart. The notion of liver zones was first described in 1954 by Rappaport^3^, who assigned hepatocytes to three zones, periportal (zone 1), midlobular (zone 2), and perivenous (zone 3), based on their distance to the nearest CV or PV. Liver zones are associated with specific functionality: lipid β-oxidation and gluconeogenesis in zone 1 and lipogenesis, ketogenesis, and glycolysis in zone 3^1^. In general, these zonated functions have been characterized with immunostaining of metabolic enzymes rather than with functional testing^1^. While individual genes were previously known to be zonated, single cell sequencing (scRNA-seq) technologies have expanded the molecular understanding of how genes and cell types are organized into zones^4–6^. To generate scRNA-seq data, cells are enzymatically dissociated so their actual spatial position can only be inferred based on gene expression landmarks. In addition, technical difficulties associated with scRNA-seq such as doublets and dropouts make these inference based methods in some ways less reliable than image-based methods. Some of these limitations could be resolved with spatial transcriptomic technologies, which would combine the geographic integrity of immunostaining with the profiling depth of scRNA-seq. With spatial approaches, there will be a critical need to measure the position of cells, features, or gene expression values with respect to zonal landmarks. These technologies are not yet available.

Despite recent advances in computer vision and deep learning, automated analysis of histology images is still immature and has not yet met the challenge of defining zonation. In the liver, applications have focused on the segmentation of cell nuclei and the prediction of ploidy states^7,8^. An unresolved problem is how to determine the position of any given modality (i.e. cell, clone, gene expression value) on an image (slide) with respect to important zonal landmarks. A key challenge is to be able to properly identify or segment zonal landmarks such as CVs or PVs, which appear as histologically similar vessel lumens lined by thin endothelial cells. Currently, approaches to distinguish these entities are based on expression of CV or PV-associated markers^9,10^. Among the studies that automatically segmented liver zones, one used geometric methods to curate histologic features and built a classifier to differentiate CVs from PVs^11^. However, the methodology was not described and was only tested on a small number of images so the performance and robustness remained uncertain. Another study used an equidistant function to divide the liver into zones^12^, however, CVs and PVs first needed to be identified manually.

Here we developed and employed a deep learning algorithm that we call the “Tissue Positioning System” (TPS), to enable quantitation of positional information, such as the zonation of liver lobules. The key rationale of TPS is that cellular features such as gene expression, proliferation, and damage, are anchored to key zonal structures in the tissue, and identification of those key structures allows for segmentation of tissue regions into biologically relevant regions. We applied TPS to liver zonation and found that it faithfully quantified the expected zonated expression patterns of the livers from 14 CreER mouse strains in an unbiased and reproducible fashion. We believe that TPS will be particularly useful for identifying tissue features that are organized into zones. In principle, the ability of TPS to define zonation around landmark structures can be adapted to any zonated tissue.

## METHODS

### Mouse strains, breeding, and slide preparation

Please refer to Wei et. al. for mouse generation and experimental details involving histology, immunohistochemistry, and immunofluorescence^13^. Briefly, mice were euthanized and liver samples were fixed in 10% Neutral buffered formalin (NBF) for 48 hours. After fixation, a liver sample was dehydrated in 30% sucrose in phosphate buffered saline (PBS) for cryosectioning. For frozen sections, slides were cut to 16 μm thickness, washed with PBS with Tween (PBST) 3x, and blocked in 5% bovine serum albumin (BSA) that contained 0.25% Triton X-100. Glutamine Synthetase (GS) antibody (Abcam ab49873) was diluted in the blocking buffer, applied to slides, and incubated overnight. Slides were imaged using an Axioscan (Zeiss) microscope and processed using Zen 2.6 software. A subset of the tissue sections and images analyzed in this paper were obtained from prior experiments shown in Wei et. al., *Science* 2021. Most of the images analyzed by TPS were independently performed for this paper. A total of 224 mice from 14 CreER lines were used in this study (**Supplementary Table S1**).

### Imaging data preparation and annotation for deep learning model development

We developed an automated annotation pipeline workflow to 1) identify vessels from the images and 2) to classify them as either CVs or PVs using information from DAPI and GS channels.

The pipeline is based on a series of image morphological operations. The first step was to segment vessels on the slide. Briefly, dark regions on the DAPI channels (DAPI masks) were identified by thresholding on a very small cutoff value (~10% of maximum DAPI intensity). Areas of each DAPI mask were calculated and the smallest ones were discarded because they were not likely to be major vascular structures (< 1% of the total image area). Lastly, because vessels were sectioned in random orientations, i.e. cross-sectioned at different angles, or even tangentially but not through the lumen, some vessels could be mistakenly removed using the vessel size cutoff. To mitigate this, we examined the very small DAPI masks and added them back if their associated GS intensity was above a GS cutoff, which was determined by applying Otsu’s thresholding^14^ on the GS channel. After these steps, the remaining DAPI masks were classified as vessel masks.

The second step was to annotate segmented vessels as CVs or PVs. We designed our pipeline based on the following features of liver lobule organization: 1) each liver lobule is centered around a CV, with surrounding PVs located on the perimeter of the lobule; 2) vessel lumens are devoid of nuclei that have high DAPI intensity; 3) CVs are lined with hepatocytes with high GS intensity.

Specifically, intensity percentile values (lower limit = 25, higher limit = 75, interval = 10) from the GS channel were quantified for each vessel mask. Then, the mask/feature data were transformed by PCA and the masks were clustered into two groups using the KMeans algorithm. Finally, the masks with higher median GS intensity were labeled as CVs, and the others as PVs.

We applied the above heuristic imaging morphological pipeline to generate the masks and classified each vessel in 182 high-quality images to develop a deep-learning based segmentation model. The classes and masks were further visually validated by pathologists. We excluded 68 images with many classification errors and obvious mask artifacts in the automatically generated masks and divided the rest of the 114 images into 61 for training, 24 for validation and 29 for testing, respectively. The labeled data was divided into 110 images for training, 36 images for validation, and 36 images for testing..

### Segmentation of CVs and PVs using deep learning

A deep-learning based segmentation algorithm was developed for CV-PV segmentation and classification. The algorithm is built with the UNet framework with the modifications described as follows. 1) We used the MobileNet-v2 backbone pre-trained on ImageNet to replace the standard CNN encoder in the original UNet. In comparison to a straightforward CNN encoder, MobileNet-v2 extracts more intricate high-level image features, and employing a pre-trained backbone boosts accuracy and lowers the risk of overfitting when the sample size is small. 2) To marginally improve performance, we swapped out the upsampling layer in the decoder for an inverted residual block plus a convolution transpose layer. The model structure is illustrated in **Figure 1b**.

**Figure 1.**
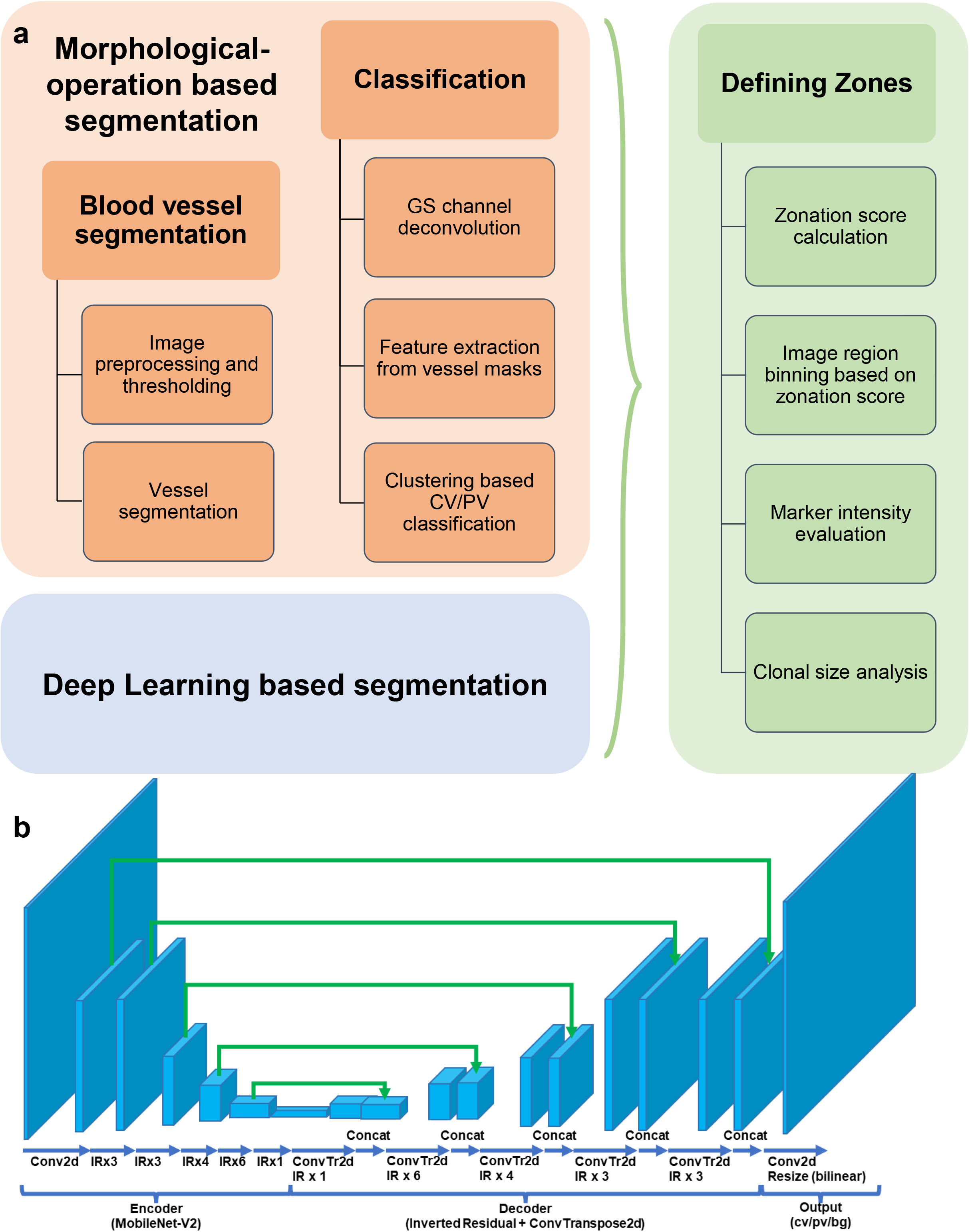
Overview of the TPS workflow.

In the training process, instead of using cross-entropy loss with pixel heatmap, soft-dice loss was used to automatically balance pixel weights for different classes. In order to prevent overfitting, color intensity augmentation, random projective transforms, and random horizontal/vertical flipping were used. In the validation process, mIoUs (median intersection-over-union) for CVs and PVs between ground truth and prediction were documented. The model with highest mIoU in the validation dataset was further evaluated on the testing dataset and deployed for downstream analysis.

### Defining position as a function of zonal landmarks

We reasoned that lobular zones were continuous in a gradient fashion along the CV-PV axis, and thus we first calculated a positional score for each pixel based on its distance to the nearest CV or PV mask. This was performed as follows for each pixel. First, the distances to the nearest CV and PV were calculated. If the pixel’s distance to the nearest CV or PV is larger than 25% of image patch size, it will be excluded from further analysis. Then, the ratio between the two distances were calculated, which reflects the position of the pixel in the context of a liver lobule. Lastly, the pixels were assigned to equal-sized groups, named TPS layers, by discretizing their distance ratios into quantile bins.

Zonal expression patterns in the tissue can be examined by evaluating marker expression in each layer. In this study, we used 24 equal-sized TPS layers. The selection of this number is largely arbitrary as the zonal expression patterns derived are not sensitive to the number of TPS layers.

### Definition of *Z_max_* and *Z_50_*

To quantitatively describe the location and selectivity of a zonal expression pattern in the context of the classic 3-zone system, we introduced two metrics, ***Z_max_*** and ***Z_50_***, respectively. ***Z_max_*** reflects the hepatocyte layer where a marker is maximally expressed. It is defined as

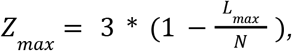

where ***L_max_*** is the layer number where a marker was expressed at maximal level, and ***N*** is the total number of TPS layers.

***Z_50_*** is the fraction of the total number of layers where marker expression is at least 50% of ***L_max_***.

It is defined as

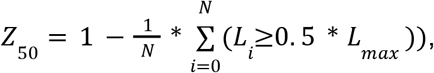

where ***L_i_*** is the expression level of marker at TPS layer ***i***.

### Measurement of clone size

The zonal expression pattern of markers can be divided into two broad categories: widely expressed in contiguous regions, or sparsely expressed in disparate regions. For sparsely expressed reporter strains, we could measure the clone number and size. Size was calculated as the marker expressing clone area or the number of nuclei in each clone. The number of nuclei in each clone was calculated using the watershed^15^ segmentation algorithm on the DAPI masks derived from Otsu^14^ thresholding.

### Analyzing whole slide images

The TPS algorithm can also analyze whole liver section images as inputs, and this was done by parallel processing of cropped images from whole liver section images. Briefly, a large input image was cropped into 4000 x 2000 images with 250-pixel margins on each edge. During each TPS segmentation iteration, a 4000 x 2000 cropped image was processed, but only results from inside the margins were kept (the central 3500 x 1500 regions). This was to ensure that vessels near the edge of the images were recognized properly. Then, the vessel masks from all cropped images were stitched together and vessel classification and expression analysis were done on the full-size image using the stitched vessel masks.

### Lobule detection

We used the watershed algorithm to detect lobules in the hepatocyte image. Briefly, distances from each pixel to the closest CV mask were calculated. Then the local maxima of the distances with each CV mask were determined. Finally, the watershed transformation was applied using the distances and local maxima to find the boundaries that separated areas around CV masks into different regions, which reflected the lobular structure in the liver.

## RESULTS

### Image-based quantification of tissue features as a function of zonal location

“Tissue Positioning System” (TPS) is software that allows automatic evaluation of tissue features as a function of zonal position within histology sections. Examples of a “tissue feature” could be any phenotypic measurement that can be assigned to a particular coordinate: i.e. gene expression, proliferation, cell size, nuclear size, clone size, lipid droplets, inflammatory cells, etc. TPS operates through two major components (see the workflow in **Figure 1**). The first component of TPS is a deep learning model that uses the nucleus channel of an image to segment CV and PV masks. In our experiment, the model could accurately identify zonal position in the 29 test slides with 0.8117 mIoU for CV and 0.8908 mIoU for PV, respectively (see **Supplementary Table S2** for performance on training, validation, and testing datasets). In the second component of TPS, positional scores proportional to the ratio between the distances of a pixel to its nearest CV and PV masks were calculated, and then pixels emanating from each concentric CV or PV were divided into equidistant bins based on this score. These bins are referred to as TPS layers. In this study, we tested the performance of TPS in determining the zonation of features such as reporter gene expression and cellular clone size.

### TPS extracts positional expression patterns from zonal reporter mice

We first tested TPS on liver sections from fluorescent reporter mice that express the Tomato fluorescent reporter in known zonal patterns. These mice (*GS-CreER*, *Cyp1a2-CreER*, and *Gls2-CreER*) were previously characterized in a qualitative fashion^13^. The accurate discernment of zone-specific reporter expression by TPS would validate the algorithm’s ability to accurately assign positional information to all pixels on an image. Each *CreER; Rosa-LSL-Tomato* mouse was given tamoxifen to activate Tomato expression prior to liver sectioning. The Tomato expression patterns for these mice are highly specific to particular zones (**Figure 2a,b**). As mentioned previously, Tomato in *GS-CreER* mice is expressed in a 1-3 hepatocyte thick ring around CVs. Tomato in *Cyp1a2-CreER* mice is expressed in all zone 3 and about 50% of zone 2 hepatocytes emanating from the CV. Tomato in *Gls2-CreER* mice is expressed in all zone 1 hepatocytes emanating from the PV. TPS identified CV and PV regions accurately without mistaking smaller gaps between cells, likely representing sinusoids, as large vessels (**Figure 2c**). To further evaluate the accuracy of predicting CVs and PVs, we predicted liver lobule boundaries associated with each CV. The lobule boundaries were very close to the center of PV masks (**Figure 2d**), indicating that the segmented CVs and PVs were reflective of true lobular structures. These results showed that TPS was able to accurately identify the landmark vessels that define liver zones in an unsupervised manner, thus providing a strong basis for the downstream assessment of tissue features.

**Figure 2.**
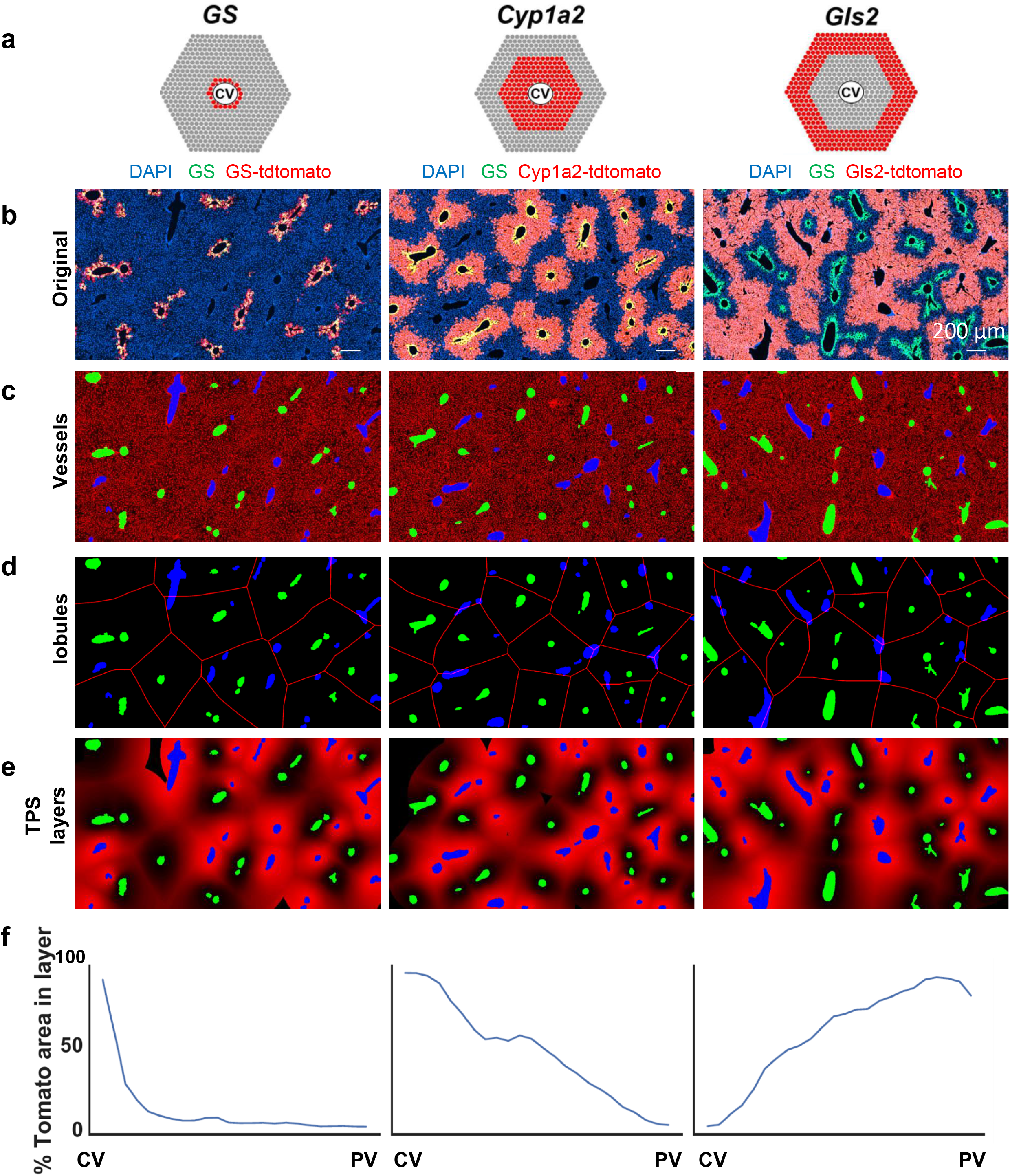
TPS characterization of Tomato expression patterns within *GS-CreER*, *Cyp1a2-CreER*, and *Gls2-CreER* mice. **a.** Schema of expected Tomato labeling in *GS-CreER*, *Cyp1a2-CreER*, and *Gls2-CreER* mice. **b.** Original three-channel immunofluorescence images of livers from CreER lineage tracing mice. Frozen sections were stained with anti-GS antibody and then scanned by a ZEISS Axio Scan.Z1 using a 20X objective (blue = DAPI, green = GS, red = Tomato). Scale bars, 100 μm. **c.** Segmented and classified CVs and PVs, shown in green and blue, respectively, for this and below panels. DAPI stains were colored red in these images. **d.** Predicted liver lobule boundaries, shown with red lines. **e.** 24 TPS layers predicted based on the pixel wise distance along the CV-PV axis. TPS layers are shown in red, and those closer to PVs are brighter. **f.** Line graphs showing the percent Tomato positive area/total area in each TPS layer. TPS layers were organized from CV to PV going from left to right. Data from one mouse is shown here.

After vessel segmentation, positional scores for all pixels were calculated and used to divide each image into 24 TPS layers reflecting zonal positions along PV-CV axes. We chose 24 for this study because at this level, zonal features of liver lobules can be adequately resolved. Pixels that were too far from any CV or PV were excluded because this likely meant that the actual nearest CV or PV was not correctly located. Consistent with lobular zonation patterns, TPS layers were organized into ring-shaped patterns around CVs and PVs (**Figure 2e**). TPS correctly showed that GS and CYP1A2 associated Tomato expression was limited to zone 3, while GLS2 associated Tomato expression was highest in zone 1 and reduced in zone 2 (**Figure 2f**). These results showed that TPS was able to reliably identify vessels, define zonal axes, and accurately quantify expression patterns in a probabilistic fashion.

### TPS discerns differences in the spatial expression patterns of genes with high resolution

Next, we evaluated TPS’s performance on each of the eleven novel *CreER* reporter mice generated by our group and the three other *CreER* mice available to the community. These fourteen zonal markers were separated into five broad categories based on their expression patterns. These include pan-zonal, all hepatocyte expressing strains (*Apoc4-CreER, Pklr-CreER*), biliary-centric strains (*Krt19-CreER*, *Sox9-CreER*), zone 1-centric strains (*Gls2-CreER, Arg1.1-CreER, Arg1.2-CreER*), zone 3-centric strains (*Axin2-CreER, Oat-CreER, Cyp1a2-CreER, GS-CreER*), and sparsely expressing strains (*Hamp2-CreER, Tert-CreER, Mup3-CreER*) (See schema in **Figure 3a**). TPS was able to accurately quantify the differences among all of these expression patterns.

**Figure 3.**
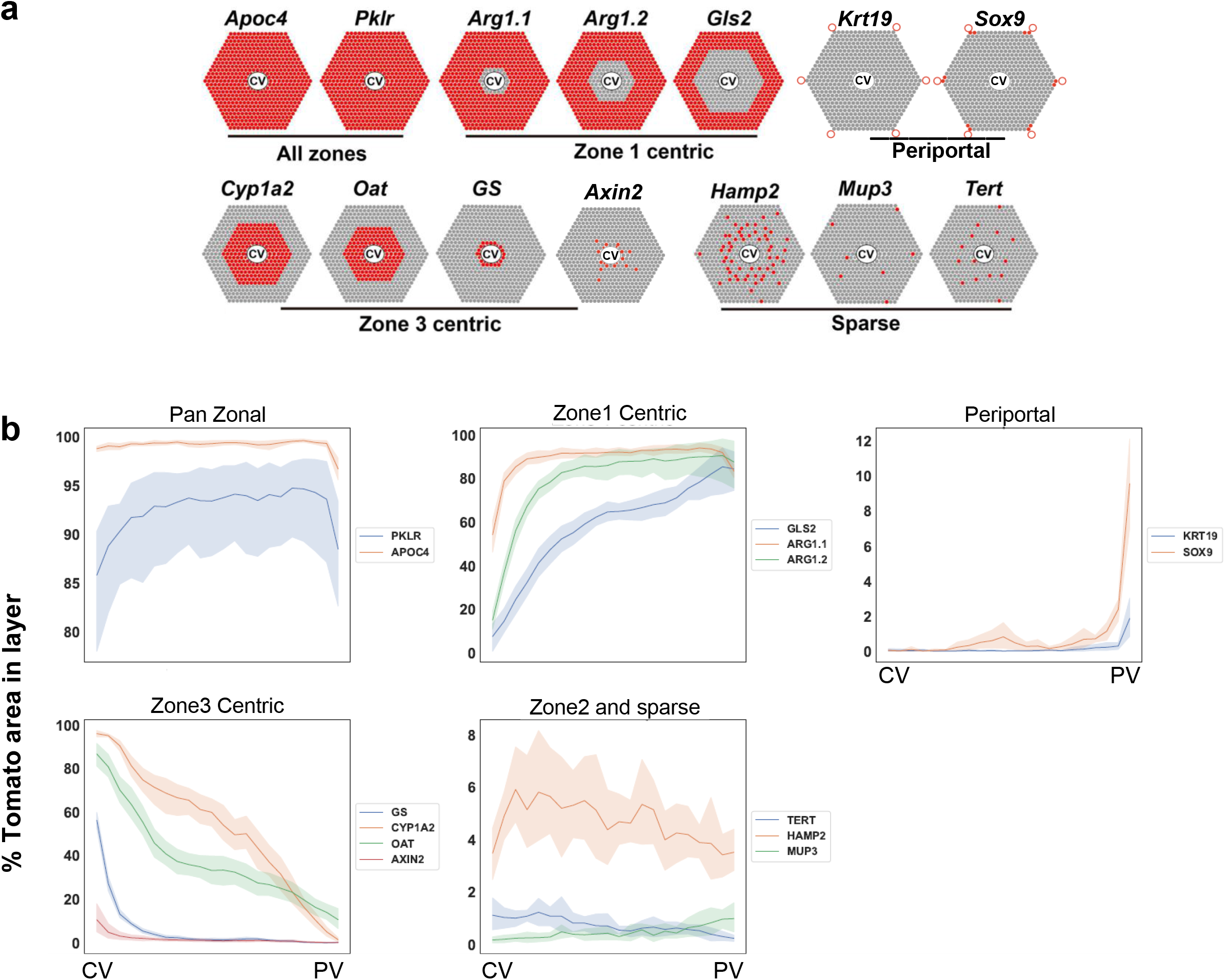
TPS is able to discern differences between qualitatively similar zonal expression patterns. **a.** Five categories of zonal expression patterns from different CreER lineage tracing mice are schematized here. All zones: *Apoc4-CreER*, *Pklr-CreER*. Zone 1 centric: *Arg1.1-CreER, Arg1.2-CreER, Gls2-CreER*. Periportal: *Krt19-CreER, Sox9-CreER*. Zone 3 centric: *Cyp1a2-CreER, Oat-CreER, GS-CreER, Axin2-CreER*. Sparse or zone 2 centric: *Hamp2-CreER, Mup3-CreER, Tert-CreER*. **b.** Line graphs showing the percentage of tomato positive area/total area in each TPS layer across the CV-PV axis. Data were collected from CreER mice 1 week after tamoxifen treatment. The 95% confidence intervals for each TPS layer are shown as shaded areas.

In addition, TPS allowed us to resolve expression patterns between reporter mice that have different expression patterns within the same zones. For example, the expression patterns of the three zone 3-centric strains (*Cyp1a2-CreER*, *Oat-CreER*, *GS-CreER*) could be clearly distinguished from one another using TPS (**Figure 3b**). To quantitatively compare zonal reporters within the same category, we introduced two metrics to characterize zonal gene expression. The zone score, **Z_max_**, describes the zonal location of peak expression, ranging from 1 to 3, corresponding to current 3-zone classification. For example, **Z_max_** of GS and GLS2 are 2.92 and 1.25, respectively. A second metric, **Z_50_**, describes the selectivity of the distribution, which measures the number of zones where a particular marker is expressed below 50% of its maximal expression level. **Z_50_** values range from 0 to 1, where 0 indicates the expression is uniform and 1 indicates the expression is restricted only to one TPS layer. For example, **Z_50_** of GS is 0.96, while **Z_50_** of Apoc4 is 0. Among all zone 3 centric markers, **Z_50_** is highest for GS and lower for Oat (0.54) and Cyp1a2 (0.46) (**Table 1**). These results showed that TPS could faithfully report zonal expression patterns and allows a more quantitative classification of expression patterns.

**Table 1.**

| **Marker** | **Z<sub>max</sub>** | **Z<sub>50</sub>** |
| --- | --- | --- |
| APOC4 | 2.5 | 0 |
| ARG1.1 | 1.5 | 0.125 |
| ARG1.2 | 1.75 | 0.21 |
| AXIN2 | 2.92 | 0.96 |
| CYP1A2 | 2.92 | 0.46 |
| GLS2 | 1.25 | 0.33 |
| GS | 2.92 | 0.96 |
| HAMP2 | 2.08 | 0.21 |
| KRT19 | 1 | 0.98 |
| MUP3 | 1 | 0.78 |
| OAT | 2.92 | 0.54 |
| PKLR | 1.92 | 0.04 |
| SOX9 | 1 | 0.96 |

### TPS is able to define the spatial patterns of sparsely expressed cells

If TPS was only able to quantitatively define the zonal expression patterns of reporter mice with obvious zone-specific expression patterns, then the algorithm would not be very useful. It would be essential for TPS to be able to determine if there is zonal specificity for features that have less obvious patterns. For instance, TPS would be useful for the characterization of reporter strains that label cells in a sparse manner because visual discernment of their zonal distribution is nearly impossible. For example, the spatial distribution of Tomato positive cells in *Hamp2-CreER* mice cannot be assessed accurately by eye because of two key reasons (**Figure S1a**). First, the area occupied by the three zones is different, and thus the total number of cells in each zone declines significantly from PV to CV. Without proper normalization for this difference in cell numbers, a perceived enrichment of labeled cells in zones that occupy larger areas would arise. Second, manual measurements are labor intensive and difficult to perform for more than a few hundred cells. For markers that express even more sparsely than Hamp2, such as Mup3 and Tert (**Figure S1b,S1c**), collecting zonal information from a few hundred cells would require analyzing dozens of large liver sections.

The first problem was solved by TPS naturally as it finds liver zones automatically. We assessed the *Hamp2-CreER* and *Mup3-CreER* reporter mice that label a small subset of hepatocytes in the liver. TPS analysis of sections from *Hamp2-CreER* mice showed that cells from any zone could express Tomato, but that midlobular zone 2 cells were labeled with the highest probability. TPS analysis of sections from *Mup3-CreER* mice also showed that cells from any zone could express Tomato, but zone 1 cells were labeled with the highest probability, followed by zones 2 and 3 (**Figure 3b**). Then we analyzed *Tert-CreER* mice created in our lab. Tert high cells were previously reported to be randomly distributed throughout the lobule based on visual inspection^10^. TPS analysis of sections from *Tert-CreER* mice showed that cells from any zone could express Tomato, but zone 3 cells were labeled with the highest probability (**Figure 3b**). TPS was able to discern the distinct patterns of each of these two reporter strains that sparsely label 0.3-10% of cells. These results show that a zonated pattern can exist even though visual inspection may only appreciate a randomly distributed pattern of expression.

For the second problem, we developed a function in TPS that can process images from full-sized liver sections in parallel, so that a large number of cells can be easily measured with analysis of whole-liver section images within a few hours on a HPC (High Performance Computer) node. We achieved this using the following steps. Each whole liver section image was divided into fixed-sized cropped images, and TPS segmentation was performed on each image in parallel (**Figure 4a,b**). Then, cropped images containing vessel masks were re-assembled and used as the vessel mask in later analyses (**Figure 4c,d**). Finally, TPS layers were defined on the original image. Zonal expression patterns derived from this large image processing were identical to those that were derived from smaller images (**Figure 4e,f**).

**Figure 4.**
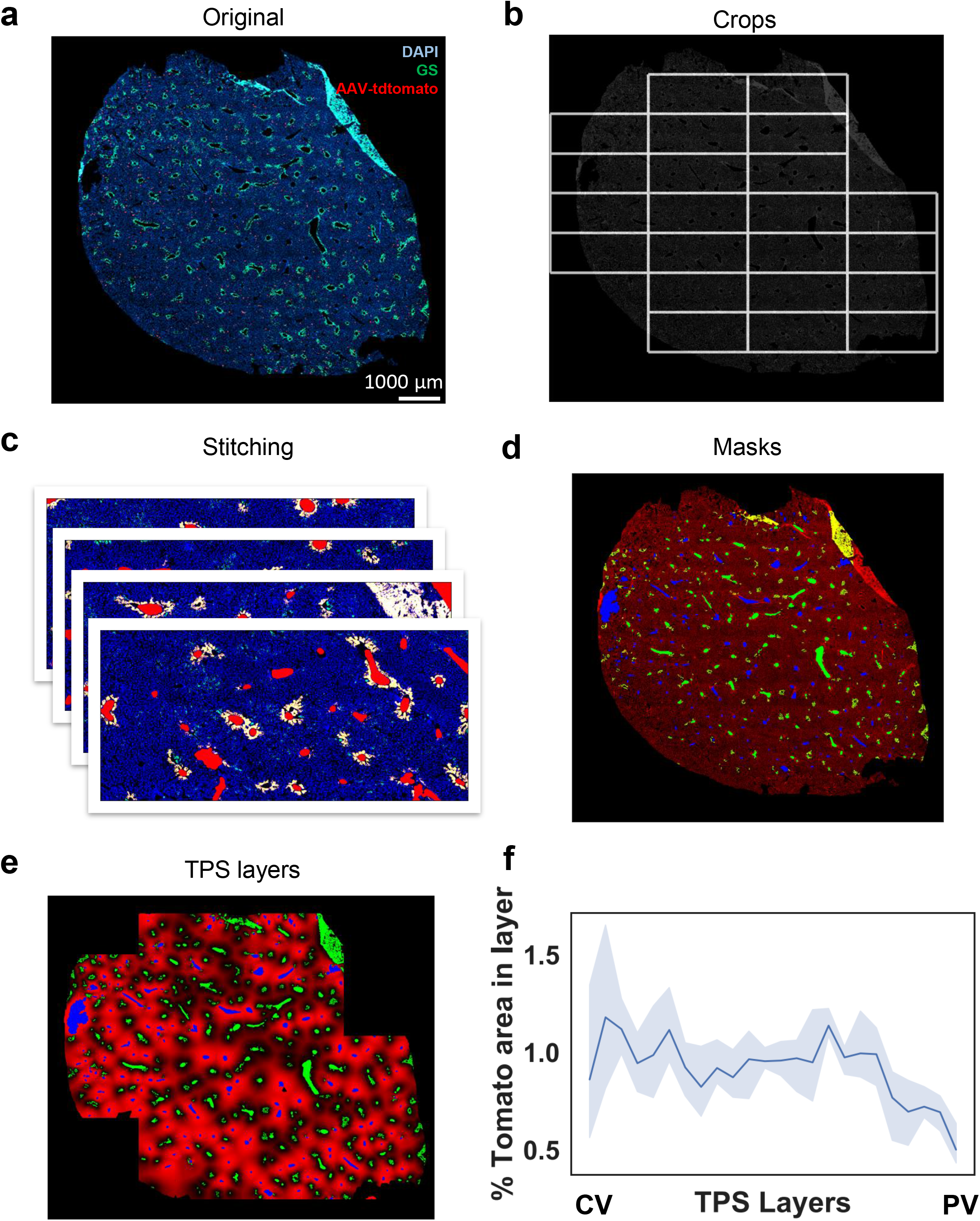
Processing of whole-liver section images by TPS. TPS can be scaled up to process whole-liver section images. This is especially useful for features that are rare or sparse, such as the low dose AAV-TBG-Cre labeling that is shown in this figure. This is achieved using a bottom up approach where each whole-liver section image is divided into cropped images and analyzed in a parallel fashion. Thus, CV/PV masks from each cropped image were pooled and used in the remaining TPS processes. **a.** Original fluorescence image obtained from Tomato reporter mice treated with low titer AAV-TBG-Cre. The liver from this AAV experiment was obtained from Wei et al and analyzed here. **b.** Cropped images used in the TPS segmentation steps. Images were extracted in a sliding window pattern, and cropped images not covering significant amounts of tissue (median DAPI intensity = 0) were not processed. Each image was set to be 3500 x 1500 pixels, but during TPS segmentation, regions of 250 pixels were added to each image to allow the correct identification of vessels near cropped image boundaries. **c.** The results from TPS segmentation on each cropped image were stitched together and used as the overall vessel mask in the following TPS processes. **d.** Shown are the CVs (green) and PVs (blue) in the stitched masks derived from all individual images. DAPI staining was colored in red. **e.** TPS layers were shown in red, and those closer to PV were brighter. CV and PV masks were shown in green and blue, respectively. **f.** Line plot of the percentage of total tomato positive area by the total cellular area in each TPS layer. The TPS layers were organized from CV to PV from left to right.

We tested this workflow using hepatocyte labeling in the *Rosa-LSL-Tomato* reporter mouse using low titer AAV-TBG-Cre injection, as is commonly used to achieve genetic recombination in a small subset of hepatocytes in a sparse but “random” fashion^16^. Indeed, TPS analysis of sections from these mice showed that hepatocytes were labeled with Tomato in an evenly distributed pattern across zones (**Figure 4f**). Next, we applied this work flow on whole-liver section images derived from *Mup3-CreER* and *Tert-CreER* reporter mice and found similar zonal expression patterns compared to those from smaller cropped images (**Figure S2**).

### TPS facilitates lineage tracing of cells during steady state homeostasis

TPS is also helpful in evaluating changes in labeling patterns during development, aging, or disease. To examine hepatocyte repopulation from different zones during steady state homeostasis, we applied TPS to all of the above *CreER* mice that were traced for over 26 weeks, or ~6 months^13^. The change in the percentage area labeled by Tomato reflects the expansion or shrinkage of domains that were labeled after tamoxifen induction. TPS analysis confirmed an expansion of cells in *Cyp1a2-CreER* and *Oat-CreER* mice, as well as a contraction of cells in *Gls2-CreER* mice (**Figure 5a**). The most significant increase occurred in zone 2 cells labeled by *Hamp2-CreER* mice. The percentage of cellular area labeled by Tomato in *Hamp2-CreER* mice increased by more than 3-fold after 6 months of tracing (from 4.63% to 14.51%; p-value < 0.0001) across all zones, but the most enriched region of expression was found near zone 2 (**Figure 5a**). Using TPS, we again found that zone 2 had a large increase in the percentage area labeled by Tomato between time points, suggesting greater expansion of cells from this midlobular zone. These results showed that TPS can extract aspects of zonal expression patterns that are very difficult to quantify using traditional methods.

**Figure 5.**
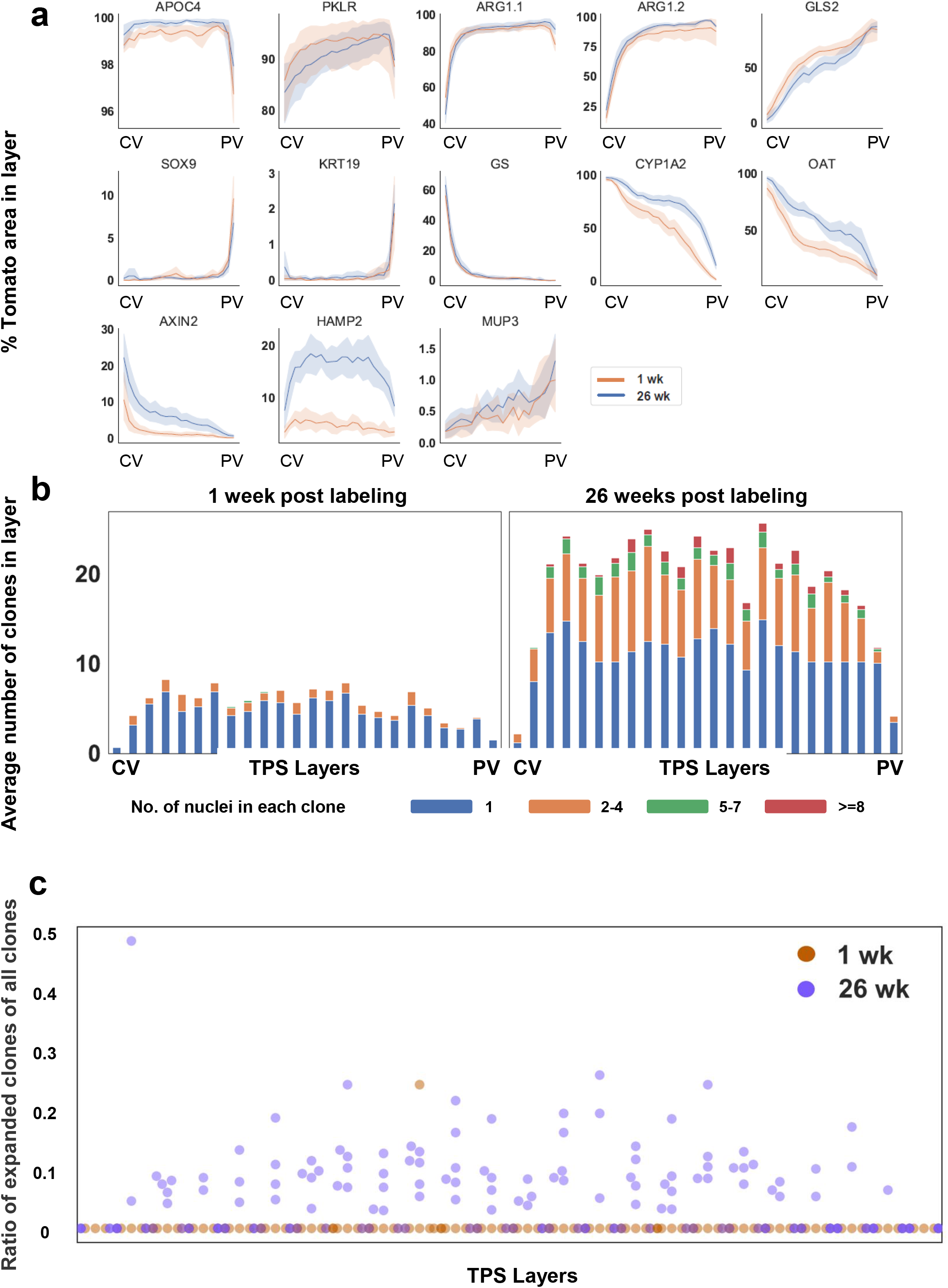
TPS revealed dynamics of liver cell subpopulations from different zones at 1 and 26 weeks after reporter labeling. TPS quantifies the changes in Tomato labeling patterns during steady state homeostasis. This analysis is able to reveal differential lineage fates of cells marked within the same zones. **a.** Comparison of lineage tracing results of 13 CreER markers 1 and 26 weeks after tamoxifen induced Tomato labeling. TPS layers were organized along the CV-PV axis. The 95% confidence interval for each TPS layer is shown as shaded areas.. **b.** Number of nuclei in each Tomato marked clone in liver sections from *Hamp2-CreER* mice. Data were grouped into four clone size categories as follows: 1 nuclei, 2-4 nuclei, 5-7 nuclei and >8 nuclei. **c.** Change in the spatial distribution of expanded clones (defined as clones with more than 3 nuclei) across hepatocyte zones in *Hamp2-CreER* mice examined at 1 and 26 weeks after tamoxifen.

### TPS is able to measure clone number, size, and location within the lobule

Beyond reporter expression, TPS was able to analyze tissue features associated with cellular clones. TPS can measure the number, size, and location of clones within lobules, information that is crucial for the understanding of regenerative activities of cells as a function of location. Here, a clone is defined as a contiguous set of reporter expressing cells surrounded by reporter negative cells. Clone size can be measured in terms of pixel area or nuclei number. The number of nuclei per clone was determined by using watershed segmentation^15^ of the DAPI channel. We identified, counted, then measured clone size in sparsely labeled reporter models such as *Hamp2-CreER* and *Mup3-CreER* mice. In *Hamp2-CreER* mice, the percentage of clones with more than 1 nuclei increased in all zones after 6 months of tracing (**Figure 5b**). This indicated that many clones expanded over time. Although expanding clones could be identified in every zone, the highest number of large clones were found in midlobular zone 2 (**Figure 5c**). Similarly, we found that Tomato labeled cells in *Mup3-CreER* mice could be found in each zone, but the largest clones (>=2 nuclei) were enriched in zone 2 (**Figure S3a,S3b**). These data confirmed that clonal expansion occurs more frequently in zone 2 compared to other zones. These results provided an example of how TPS-empowered zone segmentation can be used to systemically and reproducibly extract information about clonal dynamics as a function of location.

### TPS revealed differential dynamics from cells in different zones during injury

We also used TPS to evaluate zonal expression patterns in damaged liver tissues. To induce chronic biliary injury, we fed reporter mice with diets containing 0.1% DDC for 6 weeks, after which we harvested the liver and examined the distribution of tdTomato labeled cells. DDC is a strong cholestatic injury that models human cholangiopathies^17^. Importantly, DDC induces an inflammation related “ductular reaction” around the PV that is also seen in human liver diseases. As expected, DDC liver injury resulted in histological artifacts that complicated the analysis. For example, dark spots observed near the portal triads (**Figure 6a**) may represent cellular debris or inclusion bodies, and if coalesced together, could be mistaken as large vessels. After introducing an additional step to shrink the dark areas in the image to mitigate this possibility, TPS segmentation was able to reliably identify and distinguish CVs and PVs, and quantify Tomato expression across the lobule (**Figure 6a-c**). To examine the changes that occur in “all hepatocyte”, zone 3, and zone 1 labeled mice fed DDC, we gave tamoxifen to *ApoC4-CreER*, *GS-CreER*, and *Gls2-CreER* Tomato reporter mice, then gave them DDC food. Comparing uninjured age matched control and DDC fed livers showed that cholestatic injury caused a significant contraction of Tomato labeled zone 1 cells near the PV in both the *ApoC4-CreER* and *Gls2-CreER* strains (**Figure 6c,d**). In agreement with this, there was an increase in the number of zone 3 cells labeled with Tomato (**Figure 6c,d**). These results were consistent with the idea that DDC-induced liver damage causes loss of cells positioned in zone 1, which are in turn replaced by cells originating from zones 2 or 3. In addition, this showed that TPS was able to quantify zonal repopulation, even in the context of liver injuries that result in histological artifacts associated with tissue damage. This makes TPS a particularly useful tool for studying the cellular dynamics of livers in various settings.

**Figure 6.**
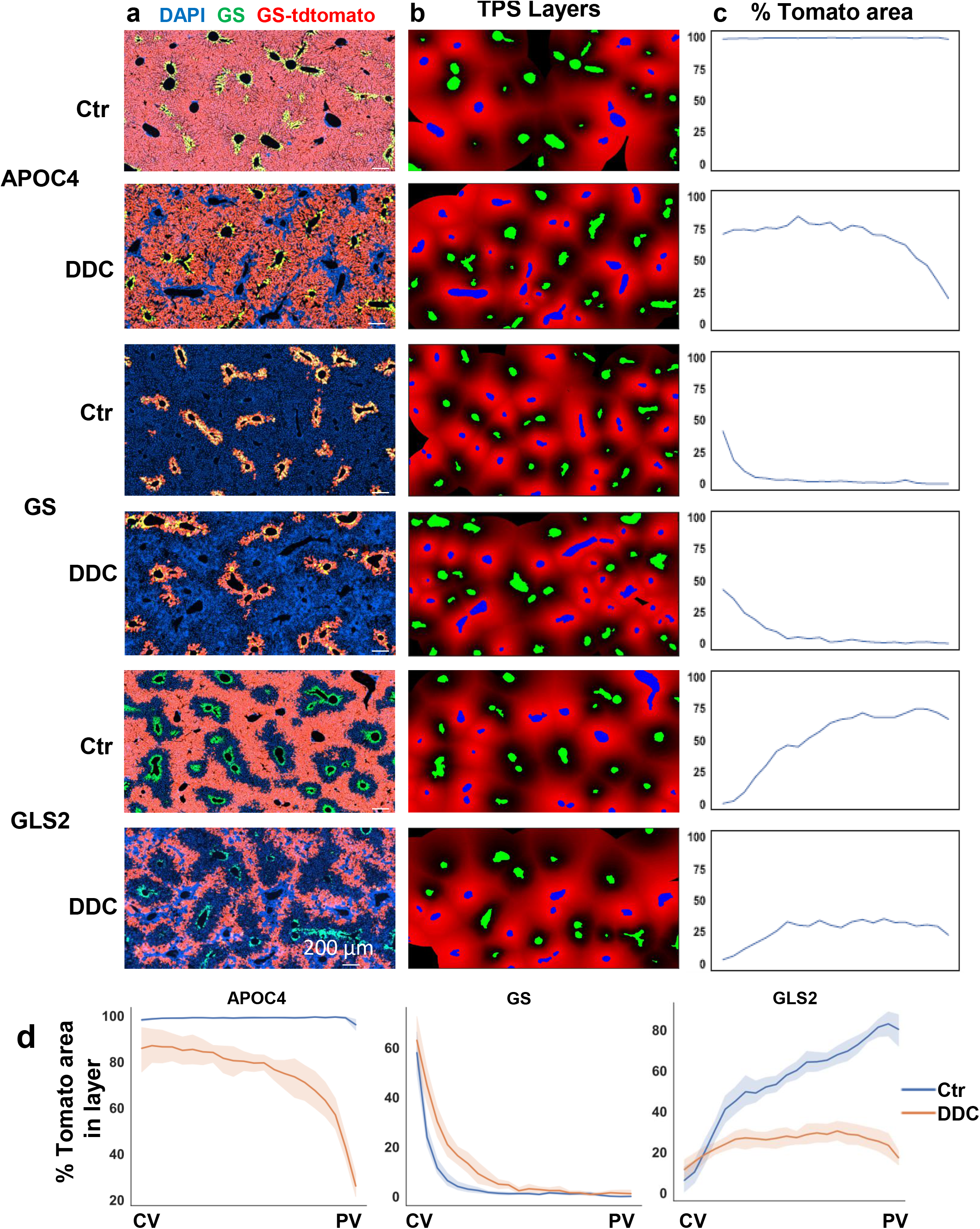
TPS revealed cellular dynamics in different zones during biliary injury. TPS is able to examine changes in Tomato labeling in livers that have been exposed to DDC. **a.** Representative original images from control and DDC treated *CreER* mice. **b.** Predicted TPS layers from each representative image. Layers were shown in red, and those closer to PV were brighter. CV and PV masks were shown in green and blue, respectively. **c.** Line plot showing the percentage of total tomato positive area by the total cellular area in each TPS layer from the representative images. The TPS layers were organized from CV to PV in left to right order. **d.** Comparison of Tomato area percentage between control and DDC treated group. The TPS layers were organized from CV to PV in left to right order.The 95% confidence interval for each TPS layer is shown as shaded areas.

## DISCUSSION

In the liver, tissue patterning or zonation is thought to be important for normal liver metabolic function^18,19^, but whether or not altered zonation might impact normal physiology and disease pathogenesis is not known. Very little is known in part because there are no computational tools to measure zonation. In addition, zonation is dynamic and can change through development and with disease, adding to the need to measure zonation over time automatically and with high precision. To our knowledge, TPS is the first fully automatic, unsupervised tool that outputs zonal information directly from input images. It can efficiently process a 4000 x 2000 image in minutes and a 20,000 x 20,000 whole tissue section image in less than half an hour.

There are several critical advantages of TPS. Currently, the identification of zonal axes requires visual analysis of images and manual discrimination of vessel identities^9,10^, which is labor intensive and susceptible to bias. TPS addresses this limitation with a deep learning based model that accurately defines CVs and PVs using only the nucleus channel of the image. This will allow high-throughput image analysis. In addition, the position of tissue features such as expression are reported as continuous variables along the CV-PV axis, rather than a qualitative measure based on the arbitrarily defined 3-zone system. This makes it possible to use more robust statistical methods to compare the zonal positions of tissue features. Because of these two advances, results from TPS are likely to be more accurate, reproducible, and comparable between different experiments.

TPS was able to determine the expression pattern of reporter mice with zone-specific expression, as well as mice with undefined, sparse expression patterns. We applied TPS to sections from *CreER* mice lines labeled with markers with different zonal expression patterns. TPS faithfully reported expected zonal expression patterns of all 14 of these reporter lines. In addition, TPS was able to quantitatively distinguish distinct strains that express reporters in a qualitatively similar fashion (i.e. within the same zone). TPS was also able to define the zonal distribution of sparsely labeled cells within lobules. The sparseness of labeling makes it difficult to determine the zonal expression pattern with manual visual analysis. For example, Tert expression in the liver was previously reported to be uniform across lobules^10^, however, TPS showed that Tert reporter expression had a modest location preference for zone 3. TPS can analyze whole liver sections, which increases the throughput and thus improves the accuracy of results. TPS is particularly suited for analyzing sparse markers to determine if there is a zone-specific expression pattern.

While we applied TPS to specific questions in liver biology, at its core TPS is able to analyze spatial location in the context of any positional landmarks. Any data associated with a position on a tissue section can be assessed with TPS. For example, any measurable tissue feature such as cell size, subcellular structures, inflammatory cells, lipid droplets, cell proliferation, or cell death can be integrated into TPS in a modular fashion. While we tested and challenged TPS with only a single gene expression readout (Tomato), it is possible to assess any number of expression values, as long as they are annotated with positional information. For example, spatial transcriptomic data, which will produce expression data for thousands of genes per cell/position, will be able to be positionally analyzed as a function of zones using TPS. Although TPS vessel segmentation is developed more specifically for the use case of liver, its ability to extract zonal expression patterns around the landmark structures in a tissue can be generalized to other organ systems and tissue types such as intestines, lungs, and lymph nodes. Thus, TPS can be adapted to other tissue types to facilitate research on biological processes that involve differential spatial patterning of any marker.

## Acknowledgements

We would like to thank Andrew Chung for constructive comments on the manuscript. T.W. is supported by CPRIT (RP190208) and NIH (CCSG 5P30CA142543). H.Z. was supported by the Pollack Foundation, NIH/NIDDK R01DK111588, and NIH/NCI R01CA251928.

## Author contributions

Y.G.W. (Yunguan Wang), T.W., G.X. and H.Z. conceived the project and wrote the manuscript.

Y.G.W. and R.R. performed the experiments and developed the TPS software.

Y.G.W. and R.R. developed the TPS algorithm and analyzed all imaging data.

Y.W. (Yonglong Wei) created the mouse models and provided images used in this manuscript.

## Conflicts

H.Z. has a sponsored research agreement with Alnylam Pharmaceuticals, consults for Flagship Pioneering and Chroma Medicines, and serves on the SAB of Ubiquitix. These interests are not directly related to the contents of this paper.

## Supplementary figures and legends

**Supplementary figure S1.**
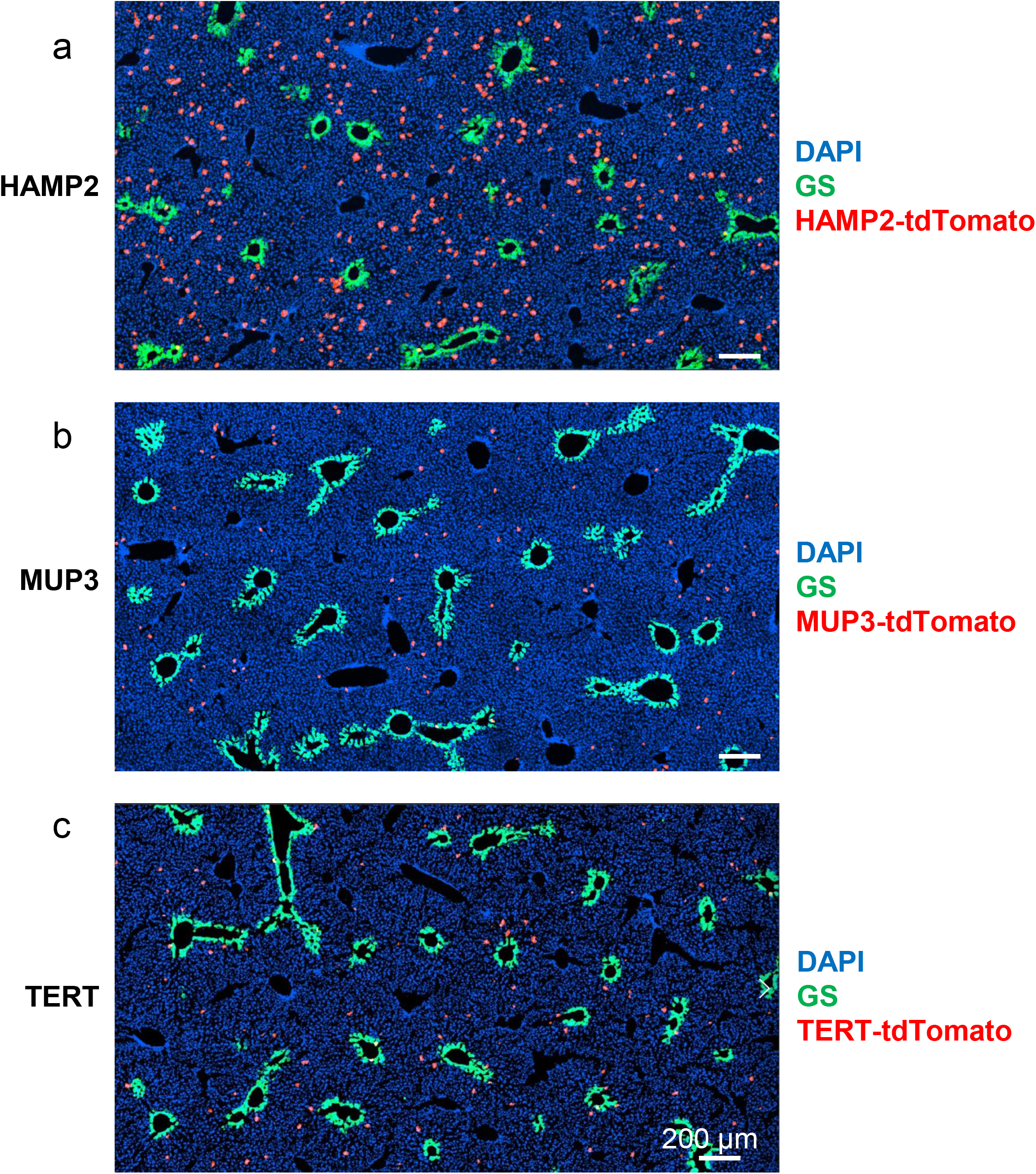
Expression patterns of *Hamp2-*, *Mup3-*, and *Tert-CreER; Rosa-LSL-tdTomato* mice. The images in this figure were obtained from liver tissue sections shown in Figure 1 of Wei et. al., Science 2021.

**Supplementary figure S2.**
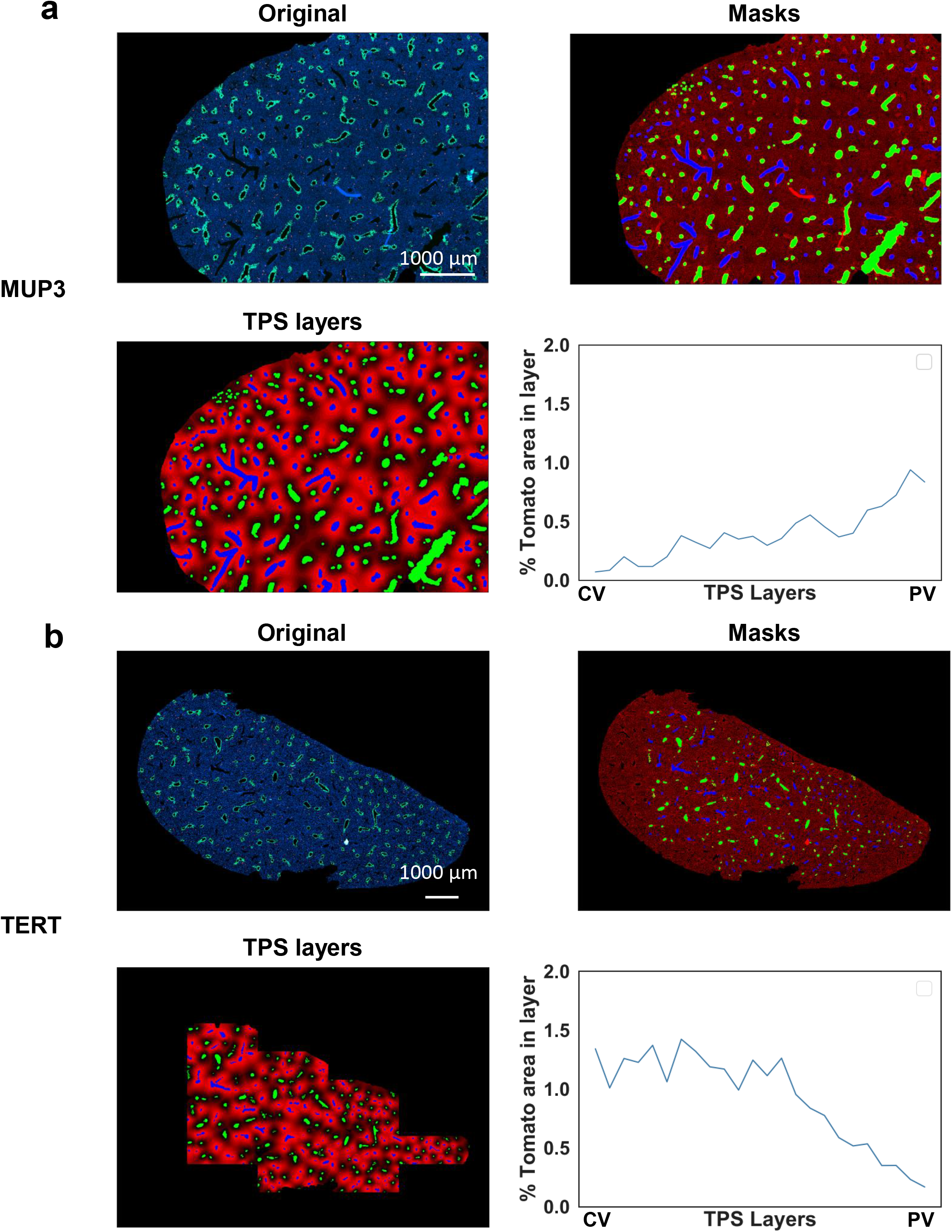
TPS analysis of whole-liver section images from *Mup3-*, and *Tert-CreER; Rosa-LSL-tdTomato* mice. Left to right, then top to bottom: original image, CV/PV masks segmented and classified by TPS, predicted TPS layers and line plot showing percentage of tomato positive area in each TPS layer. **a**. *Mup3-CreER; Rosa-LSL-tdTomato* mice and **b**. *Tert-CreER; Rosa-LSL-tdTomato* mice. The liver section in this figure was obtained from the same liver sample shown in Figure 1 of Wei et. al., Science 2021.

**Supplementary figure S3.**
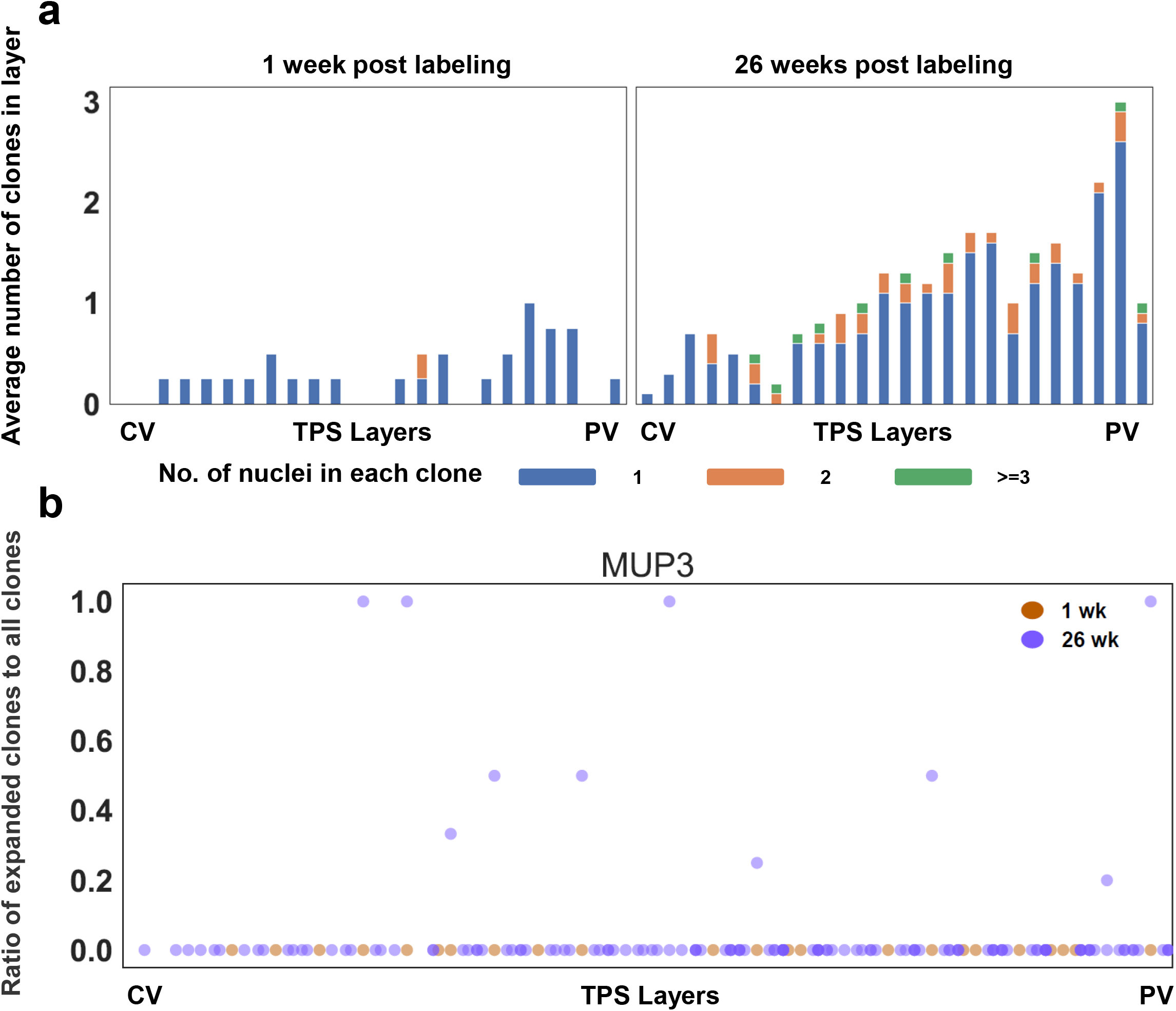
Clone size distribution in *Mup3-CreER* mice. **a.** Number of nuclei in each tomato positive clone in the liver section from *Mup3-CreER* mice labeled with Tomato. Data were grouped into 3 categories as follows: 1, 2, and >= 3 nuclei. **b.** Distribution of expanded clones (defined as clones with more than 2 nuclei) across lobular zones in *Mup3-CreER* mice 1 and 26 weeks after tamoxifen induction.

**Supplementary table S1.** Summary of the number of mice used in this paper. Summary of mice used in this study.

| **CreER strain** | **1-2 weeks** | **26 weeks** | **DDC control** | **DDC** | **Total** |
| --- | --- | --- | --- | --- | --- |
| *Apoc4-CreER* | 7 | 8 | 6 | 6 | 27 |
| *Arg1.1-CreER* | 6 | 10 |  |  | 16 |
| *Arg1.2-CreER* | 2 | 9 |  |  | 11 |
| *Axin2-CreER* | 6 | 11 |  |  | 17 |
| *Cyp1a2-CreER* | 5 | 8 |  |  | 13 |
| *Gls2-CreER* | 6 | 7 | 6 | 6 | 25 |
| *GS-CreER* | 6 | 8 | 6 | 6 | 26 |
| *Hamp2-CreER* | 6 | 7 |  |  | 13 |
| *Krt19-CreER* | 6 | 11 |  |  | 17 |
| *Mup3-CreER* | 6 | 10 |  |  | 16 |
| *Oat-CreER* | 6 | 4 |  |  | 10 |
| *Pklr-CreER* | 5 | 6 |  |  | 11 |
| *Sox9-CreER* | 6 | 10 |  |  | 16 |
| *Tert-CreER* | 6 |  |  |  | 6 |
| **Total** | **79** | **109** | **18** | **18** | **224** |

**Supplementary table S2.** Summary of TPS model performance on training, validation and test dataset. The performance of the TPS deep learning model on training, validation and testing datasets.

|  | No. image | CV mIoU | PV mIoU |
| --- | --- | --- | --- |
| Training | 61 | 0.8229 | 0.8879 |
| Validation | 24 | 0.8135 | 0.8794 |
| Test | 29 | 0.8117 | 0.8908 |

## Notes

https://github.com/yunguan-wang/Tissue_positioning_system

### Competing Interest Statement

The authors have declared no competing interest.

